# Taxonomic Description of Uncultured Cyanobacteria from Extreme Habitats through Genome-Based Classification

**DOI:** 10.64898/2026.01.02.697360

**Authors:** Edi Sudianto, Maximillian D. Shlafstein, Benoit Durieu, Marie Harmel, Luc Cornet, Jimmy H. Saw

## Abstract

Cyanobacteria form a morphologically and phylogenetically diverse group of oxygenic phototrophic bacteria inhabiting a wide range of environments, including extreme habitats such as hot springs and volcanic steam vents. Many lineages, particularly those from these extreme environments, remain uncultured and are known only from metagenome-assembled genomes (MAGs), limiting their integration into formal taxonomy. Analysis of 46 steam vent associated samples from Hawai‘i using 16S rRNA amplicon sequencing revealed that cyanobacteria dominate these communities. *Gloeobacter kilaueensis* dominated pit-like environments with low-light conditions, while Leptolyngbyaceae and other families are more dominant in structured soil and wall communities. We further reconstructed 38 high-quality cyanobacterial MAGs and incorporated them into a phylogenomic analysis comprising 343 cyanobacterial genomes, followed by genome-based comparisons against 9,026 reference genomes. This revealed eight novel species and one novel genus spanning five orders: Chroococcidiopsidales, Leptolyngbyales, Nostocales, Oculatellales, and Oscillatoriales. Following SeqCode guidelines, we provide the first formal taxonomic descriptions of cyanobacterial MAGs and propose guidelines for integrating genome-based and cultivated material. These findings highlight Hawaiian steam vents as hotspots of previously uncharacterized cyanobacterial diversity and underscore the importance of genome-based nomenclature.

## Introduction

Cyanobacteria represent a morphologically and phylogenetically diverse group of oxygenic phototrophic prokaryotes that inhabit a wide range of environments, from freshwater and marine ecosystems to terrestrial and extreme habitats such as hot springs, deserts, and polar regions (Whitton and Potts, 2012). Unlike most other prokaryotes, and due to their phototrophic nature, cyanobacteria are typically isolated and maintained as living cultures to distinguish them from potential contaminants. Accordingly, they are typically grown in a minimal medium containing only salts, minerals, and vitamin B12 (Rippka et al., 1979; Canstenholtz et al., 1981; Rippka et al., 1981; Waterbury, 2006). Weissmann et al. (2021) demonstrated that culture-based approaches are heavily biased toward copiotrophic organisms (i.e., adapted to nutrient-rich conditions), notably among cyanobacteria, while failing to capture many oligotrophs (i.e., adapted to low-nutrient environments). Recent comparisons of barcoding and genomic datasets indeed indicate that several cyanobacterial evolutionarily important lineages are absent from both cultured and genomic databases (Cornet et al., 2018; Cornet et al., 2023). Consequently, many cyanobacteria present in natural environments remain uncultured and are known only through metagenome-assembled genomes (MAGs).

These MAGs mostly remain undescribed, with no validly published taxonomic names, which limits their integration into formal microbial taxonomy and ecological analyses. Indeed, validated naming ensures clear communication among microbiologists and prevents the instability and redundancy inherent to provisional *Candidatus* names that lack official standing and nomenclatural priority (Whitman et al., 2024). The Code of Nomenclature of Prokaryotes Described from Sequence Data (SeqCode) enables the formal taxonomic description of uncultured prokaryotes based solely on genome sequences, provided that specific quality and metadata criteria are met (Hedlund et al., 2022), thereby offering an improved integration of MAGs into prokaryotic systematics. Nevertheless, the duality of cyanobacterial nomenclature—historically governed by both the International Code of Nomenclature for algae, fungi, and plants (ICN) and the International Code of Nomenclature of Prokaryotes (ICNP) (Oren, 2020; Pinevich and Averina, 2024)—complicates the taxonomic treatment of uncultured cyanobacteria. While the ICN traditionally relies on morphological descriptions and/or 16S rRNA sequences, the ICNP is primarily genome-based; consequently, integrating taxa known only from MAGs, which often lack 16S rRNA sequences (Cornet et al., 2018), across these two nomenclatural codes remains particularly challenging.

Extreme environments host microorganisms capable of surviving extreme temperatures, high salinity, intense radiation, or deep-sea pressure (Taufiq et al., 2026). In these extreme ecosystems, cyanobacteria often constitute the main primary producers (Stal, 1995; Obbels et al., 2016), yet they are notoriously difficult to cultivate. Steam vents (fumaroles) are a striking example: driven by volcanic activity, they exhibit drastic fluctuations in temperature, pH, pressure, and moisture; all of which shape their microbial communities (Hadland et al., 2024). Despite these harsh conditions, the microbial communities of Hawaiian steam vents are unexpectedly diverse (Prescott et al., 2022). Wall et al. (2015) proposed that such fumaroles act as “extremophile collectors”, serving as the biodiversity hotspot of novel extremophiles. Few cyanobacterial taxa have been isolated from Hawai‘i’s steam vents, including *Gloeobacter kilaueensis* (Saw et al., 2013) and *Fischerella* sp. JS2 and *Kovacikia* sp. JS3 (Prescott et al., 2023). Additionally, 16S rRNA sequence-based studies reveal a substantial diversity of uncultured cyanobacteria within these environments (Wall et al., 2015; Prescott et al., 2022), underscoring Hawai‘i’s steam vents as promising reservoirs of novel cyanobacterial lineages.

In this study, we analyzed 46 environmental samples collected from Hawai‘i’s steam vents and found a high relative abundance of cyanobacteria. We reconstructed 38 high-quality MAGs, among which 8 represent novel cyanobacterial species and 1 novel cyanobacterial genus, as assessed by genome-based classification methods. Specifically, we conducted phylogenomic analyses on the 38 MAGs together with a curated set of 343 reference cyanobacterial and closely related non-photosynthetic genomes and performed average nucleotide identity (ANI) and average amino-acid identity (AAI) comparisons against 9,026 cyanobacterial and melainabacterial genomes available from NCBI. We further propose the first taxonomic description of cyanobacterial MAGs in SeqCode, following the genomic criterion of ≥95% ANI for species delineation (Jain et al., 2018). Finally, we provide practical guidelines for the taxonomic description of cyanobacterial MAGs, emphasizing the integration of cultivated material.

## Materials and Methods

### Samples Collection and DNA Extraction

Detailed information regarding sampling locations and DNA extractions were described in a previous study (Prescott et al., 2022). Briefly, the samples were collected during an expedition in 2019 and came from three volcanic features on the island of Hawai‘i (also known as the Big Island) – a cave located inside the Kilauea Caldera within Hawaii Volcanoes National Park and two steam vent features located within the East Rift Zone (Prescott et al., 2022).

### Amplicon sequencing

Amplicon sequencing data targeting the V3-4 region of the 16S rRNA gene, using primers (CCTACGGGNGGCWGCAG and GACTACHVGGGTATCTAATCC), were processed using the RASPAM pipeline (Harmel et al., 2025; https://bitbucket.org/phylogeno/raspam/src/main/), in ASV mode. Analyses were performed with DADA2 v1.22.0 (Callahan et al., 2016), using the core functions *plotQualityProfile*, *filterAndTrim*, *learnErrors*, *derepFastq*, *dada*, *mergePairs*, and *assignTaxonomy*. Quality profiles were inspected to determine optimal trimming and filtering parameters, resulting in the use of truncLenF = 290, truncLenR = 240, trimLeftF = 5, trimLeftR = 5, maxEE = 2 (forward) and 5 (reverse), a minimum overlap of 12 bp, and zero mismatches allowed during merging. Chimera removal was performed using the consensus method implemented in DADA2 (Callahan et al., 2016). ASV tables were further filtered according to abundance thresholds applied both min occurrence per ASV (across samples)=100 and min occurrence of ASV in the sample=200, to retain high-confidence sequences. Taxonomic assignment of the resulting ASVs was performed using four independent reference databases: RDP trainset19 (Wang and Cole, 2024), CyanoSeq v1.3 (Lefler et al., 2023), CyanoSeq v1.3 with added species-level annotations, and CABO-16S (Eitel et al., 2025). No lineage-based filtering was applied to preserve the full taxonomic diversity. Diversity analyses and plots were generated in R v4.1.2 using the packages phyloseq v1.38.0 (McMurdie and Holmes, 2013), Biostrings v2.62.0 (Lifschitz et al., 2022), dplyr v1.1.4 (Wickham et al., 2023), vegan v2.7.1 (Dixon, 2003), and ggplot2 v4.0.0 (Wickham, 2016). Sequence identity comparison was performed using BLAST+ (Camacho et al., 2009), against a Cyanobacteria 16S reference dataset retrieved with edirect v23.9 (Kans, 2025): (esearch -db nucleotide -query “Cyanobacteria[Organism] AND 16S [Title]” | efetch -format fasta). All analyses were run using Nextflow v21.08.0 using the following commands: nextflow run main.nf --first --cpu=40 --data=’./data/2019_amplicons’ --outdir=“JIMMY_SAW” --asv; nextflow run main.nf --second --cpu=80 --data=’./data/2019_amplicons’ --outdir=“JIMMY_SAW” --asv --truncLenF=290 --truncLenR=240 --trimLeftF=5 --trimLeftR=5 --max_error_F=2 --max_error_R=5 --top_taxa=200 --feature_sample_min_reads=200 --feature_min_occurence=100 --filter_taxo=“all”

### Metagenome assembly

Raw metagenomic sequences were processed using BBDuk version 38.87 (https://sourceforge.net/projects/bbmap/) with input parameters: ktrim=r, minlen=50, mink=11, tbo=t, rcomp=f, k=21, ow=t, zl=4, qtrim=rl, trimq=20, to remove Illumina adapter sequences and low quality regions. Quality filtering excluded regions with a score less than 20 and reads under 50 bp in length. FastQC version 0.11.9 was then used to assess the quality of the trimmed sequences before downstream processing. Sequences from the same sample sites were combined before assembly to improve sequencing depth. MetaSPAdes version 3.15.4 (Nurk et al., 2017) was used to assemble the combined trimmed sequences using default parameters but with k-mers of 21, 33, 55, and 77. Seqtk version 1.3 (https://github.com/lh3/seqtk) was used to first create a list of the assembled contigs over 1kb, which were then mapped back to the trimmed reads with BBWrap version 38.87 (https://sourceforge.net/projects/bbmap/) using default parameters. For combined samples, contigs were individually mapped to each of the trimmed reads before combination. These mapped reads were sorted and indexed using Samtools version 1.10 (Danecek et al., 2021) with default parameters. The JGI summarize script that was packaged with BBmap tool was then used to determine the depths of contigs over 1,500 bp, allowing for binning with MetaBAT2 version 2.15 (Kang et al., 2019) using default parameters.

### Metagenomic Analyses

The 66 assembled MAGs were first checked for genome completeness and contamination using CheckM v1.2.2 (Parks et al., 2015) and GUNC v1.0.5 (Orakov et al., 2021) functions in the GENERA toolbox v3.0.0 (Cornet et al., 2023). Next, we selected 40 MAGs with >90% completeness and <5% contamination, meeting the data quality requirements for type genome assembly quality in the SeqCode Registry (Hedlund et al., 2022). The genomes were further classified using GTDB-Tk v2.0.0 (Parks et al., 2022) in the GENERA toolbox (Cornet et al., 2023). We run fastANI v1.33 (Jain et al., 2018) and FastAAI (Gerhardt et al., 2025) against the 9,026 Cyanobacteria/Melainabacteria genomes (7,913 cyanobacteria and 1,113 Vampirovibrionophyceae; last download date: July 10, 2025) available in the NCBI Genome Datasets (O’Leary et al., 2024) for MAG species identification based on ANI and AAI. The Hidden Markov Model profiles (pHMMs) for 115 genes associated with phycobilisomes, photosynthesis, and respiratory machinery (Lumian et al., 2021) were obtained from the KEGG database (Kanehisa et al., 2025) and queried against potential novel cyanobacterial genomes using the hmmsearch function in HMMER v3.4 (http://hmmer.org/).

To verify the phylogenetic placements of the MAGs, we combined the 38 identified cyanobacterial MAGs with 343 curated cyanobacterial and closely related non-photosynthetic genomes from Sudianto et al. (2025). A supermatrix was generated for the 381 taxa using GToTree v1.8.8 (Lee, 2019) with 251 individual alignments produced using the cyanobacterial pHMMs available in the tool. Sequence alignment and trimming were performed with MUSCLE v5.1 (Edgar, 2022) and TrimAl v1.4.rev15 (Capella-Gutiérrez et al., 2009), respectively. GToTree was run with minor parameter adjustments (“-c 1 -G 0”). We used IQ-TREE v2.3.6 (Minh et al., 2020), with automated ModelFinder for model selection (-m MFP; Kalyaanamoorthy et al., 2017) and 1,000 ultrafast bootstrap replicates (Hoang et al., 2018), to infer the phylogenomic tree. The resulting tree and associated information (e.g., genome completeness, contamination, ANI%) were visualized using iTOL v7 (Letunic and Bork, 2024).

## Results

### Amplicon sequencing

From 10,879,048 raw reads, 4,424,992 passed filtering, 4,005,210 merged successfully, and 2,438,362 non-chimeric reads were retained, yielding 18,830 ASVs. Abundance-based filtering kept ASVs present in ≥200 reads per sample and occurring ≥100 times across the dataset, resulting in 2,542 high-confidence ASVs. The four sampling sites harbor distinct bacterial communities, largely dominated by Cyanobacteriota and Chloroflexota (**Figure 1**). In the open area of the East Rift Zone, near the pit-like feature, Cyanobacteriota represent 49.2% of the community and are dominated by Hapalosiphonaceae (64.5%), Scytonemataceae (14.9%), and Oculatellaceae (11%) (**Table S1**). These communities contrast with those found inside the pit-like feature, where Cyanobacteriota reach 79% and are strongly dominated by the family Gloeobacteraceae (85.2%). Inside the Big Ell cave (Kīlauea Caldera), community composition varies between the soil—dominated by Chloroflexota (60%)—and the wall, which harbors a more balanced assemblage of Cyanobacteriota (47.7%) and Chloroflexota (40%). We further identified five rare cyanobacterial ASVs with BLAST sequence similarity ranging from 86.23–93.15% (**Table 1**). Following CyanoSeq classification, the lineage of these five ASVs is either Prochlorococcaceae (3), Leptolyngbyaceae (1), or broadly Cyanophyceae (1). Interestingly, these rare cyanobacterial ASVs are found only in pit-like areas, suggesting they may be the source of atypical or novel cyanobacterial spots.

**Figure 1:**
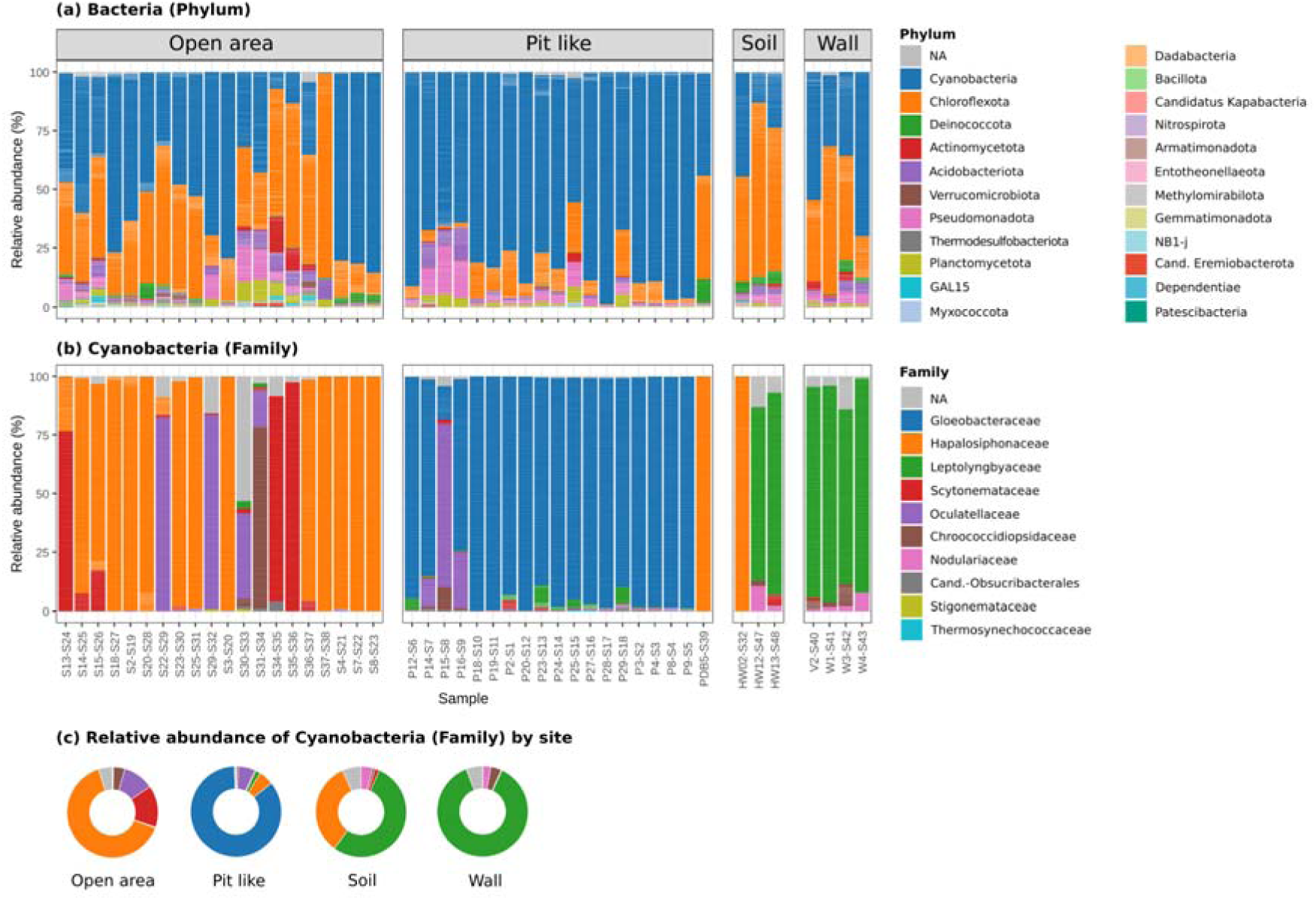
Abundance of cyanobacteria from 16S rRNA Amplicon Sequencing. Description of the bacterial diversity found on the four sampling sites. **(a)** Relative abundance of Cyanobacteria compared to other phylums of bacteria, **(b)** relative abundance of Family within Cyanobacteria, **(c)** relative abundance of Cyanobacterial family by site.

**Table 1.** Rare cyanobacterial ASV sequences from steam vents.

| ASV Name | BLAST Searches |  |  | CyanoSeqV1.3 lineage | No. of detected samples at sampling sites |  |  |  |  |  |  |
| --- | --- | --- | --- | --- | --- | --- | --- | --- | --- | --- | --- |
|  | Hit ID | Description | Sequence Similarity (%) |  | P14 | P16 | P24 | P25 | P27 | P28 | P29 |
| ASV_1789 | OR727807.1 | Unicellular cyanobacterium A47 | 93.151 | Bacteria;Cyanobacteriota;Cyanophyceae; Synechococcales;Prochlorococcaceae;Syn echococcus_B | 0 | 0 | 0 | 0 | 0 | 0 | 135 |
| ASV_2010 | AM940917.1 | Uncultured cyanobacterium partial 16S rRNA gene; clone B9_86 | 93.072 | Bacteria;Cyanobacteriota;Cyanophyceae; Leptolyngbyales;Leptolyngbyaceae | 61 | 0 | 53 | 0 | 0 | 6 | 0 |
| ASV_2139 | AM940917.1 | Uncultured cyanobacterium partial 16S rRNA gene; clone B9_86 | 93.023 | Bacteria;Cyanobacteriota;Cyanophyceae | 0 | 86 | 0 | 20 | 0 | 8 | 0 |
| ASV_2222 | KC262697.1 | Uncultured cyanobacterium clone OTUC33 | 86.404 | Bacteria;Cyanobacteriota;Cyanophyceae; Synechococcales;Prochlorococcaceae;Syn echococcus_B | 0 | 0 | 0 | 0 | 111 | 0 | 0 |
| ASV_2472 | GU271464.1 | Uncultured Trichodesmium sp. clone 1_241 | 86.23 | Bacteria;Cyanobacteriota;Cyanophyceae; Synechococcales;Prochlorococcaceae;Syn echococcus_B | 0 | 0 | 0 | 0 | 0 | 0 | 102 |

### Identification of novel cyanobacterial MAGs

Out of the 66 cyanobacterial MAGs, 40 MAGs with high CheckM completeness (>95%) and low contamination scores (<5%) were selected. These MAG sizes range from 4.5–7.9 Mb and ANI of 78.7–99.9% to available cyanobacterial genomes (**Table S2**). Two MAGs (S29-31_bin.128 and P18-20_bin.4) that are potentially Vampirovibrionaphyceae were excluded from further analyses, resulting in 38 cyanobacterial MAGs. Phylogenomic tree indicates that these 38 MAGs are nested within the Gloeobacterales, Oculatellales, Leptolyngbyales, Oscillatoriales, Chroococidiopsiales, and Nostocales (**Figure 2**). Twenty two of these MAGs can be identified down to the species level (ANI≥95%; **Figure 2**; **Table S2**), such as *Gloeobacter kilaueensis* (P23-25_bin.50, P27-29_bin.2, P18-20_bin.17, P2-4_bin.3, and P8-12_bin.7), cyanobacteriota bacterium JS3 (W1-4_bin.70^TS^, HW-13_bin.4, P23-25_bin.53, and V2_bin.1), *Fischerella* sp. FACHB-330 (S18-20_bin.32, S34-36_bin.84, S13-15_bin.40, and S2-4_bin.62), and cyanobacterium PCC 7702 (HW-02_bin.18, S2-4_bin.85, S34-36_bin.68, PDB5_bin.16, S13-15_bin.112, S22-25_bin.34, S7-8_bin.7, and S18-20_bin.8). The 16 remaining MAGs have ANI<95%, indicating the genomes of potential new cyanobacteria. Here, we describe nine novel cyanobacterial genomes according to the SeqCode’s classification, comprising eight novel species and one novel genus (**Table 2**). The AAI scores are >65% to the closest cyanobacteria genome available in NCBI for these nine MAGs (**Table S2**). Nevertheless, the phylogenetic placement of W1-4_bin.36^TS^ between the Nostocaceae and the Scytonemataceae (**Figure 2**) indicates the presence of a novel genus.

**Figure 2:**
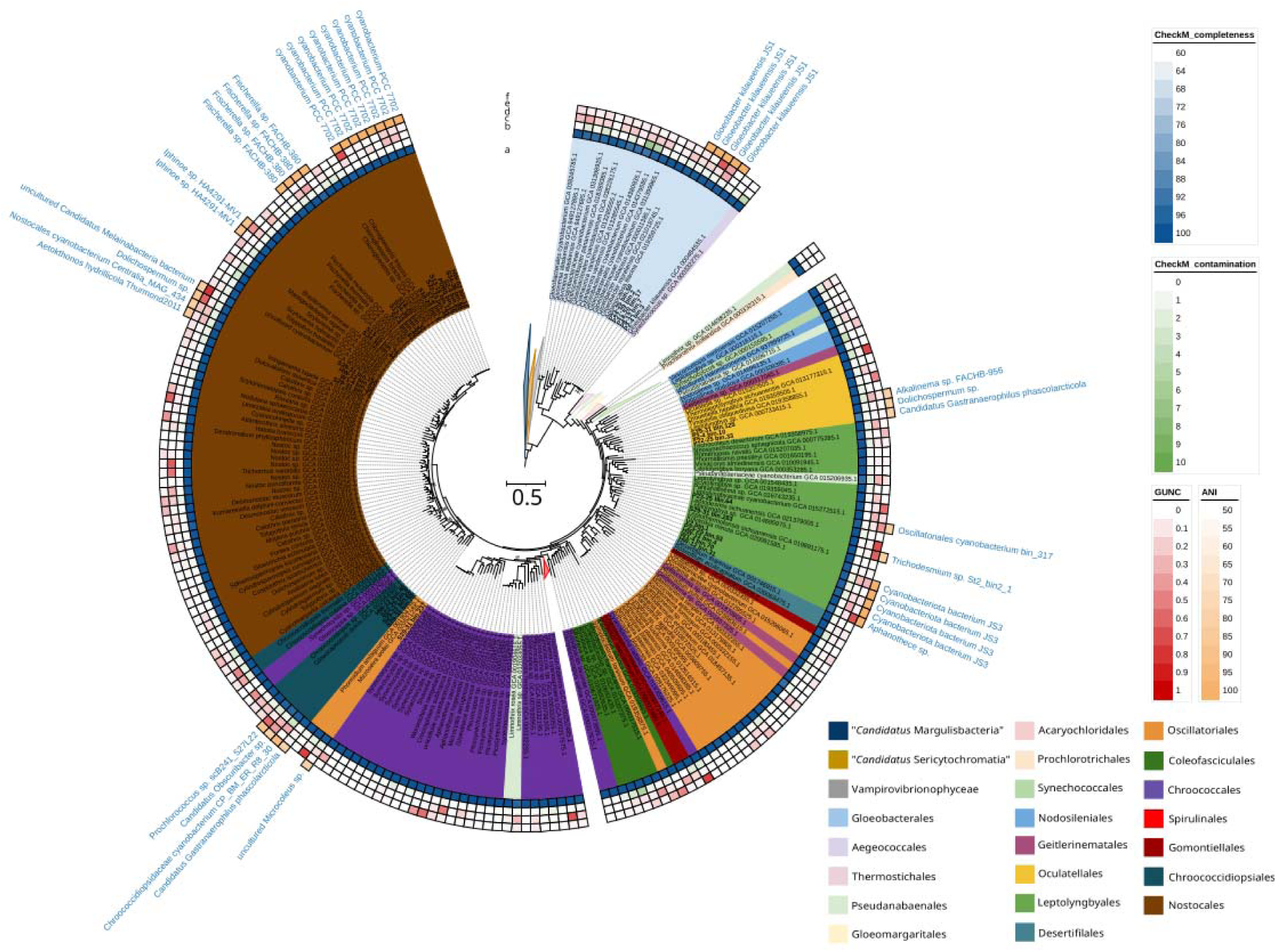
Maximum likelihood phylogenomic tree of 381 cyanobacterial genomes, including newly sequenced genomes labeled in bold. The tree was inferred from 46,754 unambiguously aligned sites under the best-fit model (LG+F+I+R10) and 1,000 ultrafast bootstrap replicates. Only nodes with <90% bootstrap supports are labeled. Annotations include: (a) cyanobacterial order assignments following Strunecký et al. (2023); (b) CheckM completeness; (c) CheckM contamination; (d) GUNC CSS; (e) GUNC contamination; (f) Percentage ANI and best-match taxa.

**Table 2:** Statistics and applied guidelines of selected cyanobacterial MAGs deposited in SeqCode.

| MAG | Accession No. | Genome size (Mb) | No. of contigs | Completeness (%) | Contamination (%) | GTDB Classification | ANI (%) | Proposed names | Guidelines applied* |
| --- | --- | --- | --- | --- | --- | --- | --- | --- | --- |
| W1-4_bin.70<sup>TS</sup> | ERZ28674871<sup>TS</sup> | 6.30 | 227 | 98.82 | 0.47 | s__Kovacikia sp034541955 | 99.6772 | *Kovacikia ellensis* | A3: Description and recommend deposit |
| W1-4_bin.36<sup>TS</sup> | ERZ28674878<sup>TS</sup> | 7.88 | 195 | 99.04 | 1.59 | s__JAJPJF01 sp021324415 | 81.405 | *Paranostoc speluncae* | B4: < 95 % ANI |
| S29-31_bin.74<sup>TS</sup> | ERZ28674877<sup>TS</sup> | 7.66 | 587 | 96.75 | 1.81 | s__Aetokthonos hydrillicola | 85.3098 | *Aetokthonos rifticola* | B4: < 95 % ANI |
| S29-31_bin.60<sup>TS</sup> | ERZ28674876<sup>TS</sup> | 6.44 | 173 | 97.56 | 3.22 | s__JAF AVL01 sp019244235 | 94.3411 | *Chroococcidiopsis rifticola* | B4: < 95 % ANI |
| HW-12_bin.40<sup>TS</sup> | ERZ28674869<sup>TS</sup> | 5.72 | 367 | 96.89 | 2.23 | s__Prochlorococcus_A sp001989455 | 86.9307 | *Chroococcidiopsis edaphicus* | B4: < 95 % ANI |
| S29-31_bin.237<sup>TS</sup> | ERZ28674875<sup>TS</sup> | 7.83 | 105 | 98.67 | 1.56 | s__JANRIL01 sp033764955 | 79.1589 | *Microcoleus pahoensis* | B4: < 95 % ANI |
| S29-31_bin.125<sup>TS</sup> | ERZ28674874<sup>TS</sup> | 6.86 | 283 | 98.82 | 2.2 | s__MUGG01 sp014697025 | 79.9028 | *Elainella pahoensis* | B4: < 95 % ANI |
| S13-15_bin.75<sup>TS</sup> | ERZ28674873<sup>TS</sup> | 6.81 | 326 | 98.92 | 0.44 | s__Brasilonema sp019359415 | 89.9154 | *Brasilonema rifticola* | B4: < 95 % ANI |
| P23-25_bin.64<sup>TS</sup> | ERZ28674872<sup>TS</sup> | 7.54 | 213 | 97.41 | 0.71 | s__Leptodesmis sp037442585 | 79.7857 | *Leptodesmis pahoensis* | B4: < 95 % ANI |

\*The applied guidelines followed the classification in Figure 3

### Photosynthetic potential of the novel cyanobacterial MAGs

As oxygenic photosynthesis is the synapomorphy of cyanobacteria, we used 115 genes associated with the phycobilisome, photosynthetic apparatus, and respiratory machinery to characterize the novel cyanobacterial species identified from our MAGs (**Table S3**). W1-4_bin.70^TS^ encodes the fewest, largely because it lacks the cytochrome bd complex (*cydABX*). In contrast, the genomes of S29-31_bin.237^TS^ and W1-4_bin.36^TS^ each encode 100 of the 115 genes. All MAGs encode a complete or nearly complete phycobilisome gene set. The core allophycocyanin genes (*apcABCDEF*) and rod phycocyanin/phycoerythrocyanin genes (*cpcABCDEFG*) are present in all MAGs. The phycoerythrin gene set (*cpeABCDERSTUYZ*) is broadly conserved; however, the *cpeR* gene is absent from five out of the nine MAGs, specifically P23-25_bin.64^TS^, S29-31_bin.125^TS^, S29-31_bin.237^TS^, S29-31_bin.60^TS^, and W1-4_bin.70^TS^. Several photosystem I and II genes (e.g., *psaGHNO* and *psbQRW*) were absent from these nine cyanobacterial MAGs as they are eukaryote-specific subunits. All nine MAGs contain a full complement of 29 genes associated with F-type ATPase, NADH dehydrogenase, and photosynthetic electron transport chains. The only gene missing from any genome is *hoxE,* which is absent from S13-15_bin.75^TS^, S29-31_bin.125^TS^, and W1-4_bin.70^TS^ (**Table S3**). Several genes responsible for the cytochrome b6f complex (*petGN*), cytochrome c oxidase (*ctaF*), and succinate dehydrogenase (*sdhCD*) are missing from most of the nine MAGs. Specifically, *petG* occurs in 2/9 MAGs and *petN* in 1/9, whereas *ctaF*, *sdhC*, and *sdhD* are not detected in any MAG (**Table S3**).

### SeqCode description

#### Description of Kovacikia ellensis sp. nov

*Kovacikia ellensis* (el’len.sis (N.L. masc./fem. adj. *ellensis*), from Ell-, referring to the “Big Ell” cave; L. suffix -*ensis*, meaning originating from or pertaining to; N.L. masc./fem. adj. *ellensis*, originating from the Ell cave). Four MAGs corresponding to this species were derived from samples collected from diverse sites (open area, wall, soil) of the Big Ell cave, Hawai‘i. The type genome (W1-4_bin.70^TS^) is 6.30 Mb, with CheckM completeness of 98.82% and contamination of 0.47%. This species has been previously cultured and has its genome deposited as “cyanobacteriota bacterium JS3” in the NCBI (GCA_034541955.1, ANI: 99.6%; AAI: 82.8%) by Prescott et al. (2023), but has not been officially named using any nomenclature guidelines. Here, we proposed using the SeqCode nomenclature to name this species. The genome lacks a complete cytochrome bd complex (*cydABX*), but retains genes related to the cytochrome b6f complex (*petABCDLM*) and cytochrome c oxidase (*ctaCDE*). In total, this MAG encodes 96 (out of 115) genes related to phycobilisome, photosynthesis, and respiratory machinery. The type genome has been deposited in ENA with the accession number ERZ28674871^TS^. Phylogenomic analysis placed this genome in the order Leptolyngbyales. *Kovacikia ellensis* belongs to the family Leptolyngbyaceae, order Leptolyngbyales, class Cyanophyceae, and phylum Cyanobacteriota. The name has been registered under SeqCode: seqco.de/r:s5njm2s9.

#### Description of Paranostoc speluncae sp. nov

*Paranostoc speluncae* (spe.lun’cae (N.L. gen. n./adj. *speluncae*), L. fem. n. *spelunca*, cave; L. gen. sg. ending -*ae*, indicating association or inhabitation; N.L. gen. n./adj. *speluncae*, inhabiting or associated with a cave). This MAG (W1-4_bin.36^TS^) has a size of 7.8 Mb, completeness of 99.04%, and contamination of 1.59%. This MAG was assembled from sequencing reads generated from samples taken from a cave wall located near the entrance of Big Ell. This is the same wall from which *Gloeobacter kilaueensis* JS1 was isolated in 2013 (Saw et al., 2013). The closest identified species is a Nostocales cyanobacterium (GCA_021324415.1, ANI: 81.4%; AAI: 74.3%). In total, this MAG encodes 100 (out of 115) genes related to phycobilisome, photosynthesis, and respiratory machinery. This type genome has been deposited in ENA with the accession number ERZ28674878^TS^. Phylogenomic analysis placed this genome in the order Nostocales. *Paranostoc* belongs to the family Stigonemataceae, order Nostocales, class Cyanophyceae, and phylum Cyanobacteriota. The name has been registered under SeqCode: seqco.de/r:s5njm2s9.

#### Description of Aetokthonos rifticola sp. nov

*Aetokthonos rifiticola* (rif.ti’co.la (N.L. masc. n. *rifticola*), N.L. neut. n. *riftum*, “rift”; derived from English rift; L. n. -*cola*, an inhabitant or dweller; N.L. masc. n. *rifticola*, one that dwells in a rift). This MAG (S29-31_bin.74^TS^) has a size of 7.6 Mb, completeness of 96.75%, and contamination of 1.81%. This MAG was assembled from sequencing reads generated from samples taken from an open area located within a publicly accessible steam vent features located within the East Rift Zone. The closest identified species is *Aetokthonos hydrillicola* (GCA_017591595.2, ANI: 83.9%; AAI: 79.5%). In total, this MAG encodes 99 (out of 115) genes related to phycobilisome, photosynthesis, and respiratory machinery. This type genome has been deposited in ENA with the accession number ERZ28674877^TS^. Phylogenomic analysis placed this genome in the order Nostocales. *Aetokthonos rifiticola* belongs to the family Scytonemataceae, order Nostocales, class Cyanophyceae, and phylum Cyanobacteriota. The name has been registered under SeqCode: seqco.de/r:s5njm2s9.

#### Description of Chroococcidiopsis rifticola sp. nov

*Chroococcidiopsis rifticola* (rif.ti’co.la (N.L. masc. n. *rifticola*), N.L. neut. n. *riftum*, “rift”; derived from English rift; L. n. -*cola*, an inhabitant or dweller; N.L. masc. n. *rifticola*, one that dwells in a rift). This MAG (S29-31_bin.60^TS^) was assembled from sequencing reads generated from samples taken from an open area located within a publicly accessible steam vent features located within the East Rift Zone. S29-31_bin.60^TS^ has a genome size of 6.4 Mb, completeness of 97.6%, and contamination of 3.2%. Both ANI and AAI identified it as a Chroococcidiopsidaceae cyanobacterium (GCA_019244235.1, ANI: 94.3%; GCA_019242465.1, AAI: 76.9%). In total, this MAG encodes 99 (out of 115) genes related to phycobilisome, photosynthesis, and respiratory machinery. This type genome has been deposited in ENA with the accession number ERZ28674876^TS^. Phylogenomic analysis placed this genome in the order Chroococcidiopsidales. *Chroococcidiopsis rifticola* belongs to the family Chroococcidiopsidaceae, order Chroococcidiopsidales, class Cyanophyceae, and phylum Cyanobacteriota. The name has been registered under SeqCode: seqco.de/r:s5njm2s9.

#### Description of Chroococcidiopsis edaphicus sp. nov

*Chroococcidiopsis edaphicus* (ed.a’phi.cus (N.L. masc. adj. *edaphicus*), Gr. neut. n. [δαφος *edaphos*, “soil, ground”; N.L. adj. suffix -*icus* (from Gr. -ικός, “pertaining to”); N.L. masc. adj. *edaphicus*, pertaining to soil). This MAG (HW-12_bin.40^TS^) was assembled from sequencing reads generated from samples taken from a soil from Big Ell, a cave located within Kilauea Caldera inside Hawaii Volcanoes National Park. HW-12_bin.40^TS^ has a genome size of 5.7 Mb, completeness of 96.9%, and contamination of 2.2%. The closest species identification was *Prochlorococcus* (GCA_000635355.1, ANI: 86.9%) and *Chroococcidiopsis* sp. CCMEE29 (GCA_023558375.1, AAI: 65.9%). In total, this MAG encodes 98 (out of 115) genes related to phycobilisome, photosynthesis, and respiratory machinery. This type genome has been deposited in ENA with the accession number ERZ28674869^TS^. Phylogenomic analysis placed this genome in the order Chroococcidiopsidales. *Chroococcidiopsis edaphicus* belongs to the family Chroococcidiopsidaceae, order Chroococcidiopsidales, class Cyanophyceae, and phylum Cyanobacteriota. The name has been registered under SeqCode: seqco.de/r:s5njm2s9.

#### Description of Microcoleus pahoaensis sp. nov

*Microcoleus pahoaensis* (pa.ho.a.en’sis (N.L. adj. *pahoaensis*), from Pāhoa, a toponym in Hawai‘i; L. suffix -*ensis*, pertaining to or originating from; N.L. adj. *pahoaensis*, originating from Pāhoa). This MAG (S29-31_bin.237^TS^) has a size of 7.8 Mb, completeness of 98.67%, and contamination of 1.56%. This MAG was assembled from sequencing reads generated from samples taken from an open area located within a publicly accessible steam vent features located within the East Rift Zone. The closest identified species is a Thermostichaceae (GCA_033764955.1, ANI: 79.1%) or an uncultured *Microcoleus* (GCA_964657555.1, AAI: 71.1%). In total, this MAG encodes 100 (out of 115) genes related to phycobilisome, photosynthesis, and respiratory machinery. This type genome has been deposited in ENA with the accession number ERZ28674875^TS^. Phylogenomic analysis placed this genome in the order Oscillatoriales. *Microcoelus pahoaensis* belongs to the family Microcoleaceae, order Oscillatoriales, class Cyanophyceae, and phylum Cyanobacteriota. The name has been registered under SeqCode: seqco.de/r:s5njm2s9.

#### Description of Elainella pahoaensis sp. nov

*Elainella pahoaensis* (pa.ho.a.en’sis (N.L. adj. *pahoaensis*), from Pāhoa, a toponym in Hawai‘i; L. suffix -*ensis*, pertaining to or originating from; N.L. adj. *pahoaensis*, originating from Pāhoa). This MAG (S29-31_bin.125^TS^) has a size of 6.8 Mb, completeness of 98.82%, and contamination of 2.2%. This MAG was assembled from sequencing reads generated from samples taken from an open area located within a publicly accessible steam vent features located within the East Rift Zone. The closest identified species is an *Alkalinema* (GCA_014697025.1, ANI: 79.9%) or *Elainella* (GCA_000733415.1, AAI: 74.3%). In total, this MAG encodes 97 (out of 115) genes related to phycobilisome, photosynthesis, and respiratory machinery. This type genome has been deposited in ENA with the accession number ERZ28674874^TS^. Phylogenomic analysis placed this genome in the order Oculatellales. *Elainella pahoaensis* belongs to the family Oculatellaceae, order Oculatellales, class Cyanophyceae, and phylum Cyanobacteriota. The name has been registered under SeqCode: seqco.de/r:s5njm2s9.

#### Description of Brasilonema rifticola sp. nov

*Brasilonema rifticola* (rif.ti’co.la (N.L. masc. n. *rifticola*), N.L. neut. n. *riftum*, “rift”; derived from English rift; L. n. -*cola*, an inhabitant or dweller; N.L. masc. n. *rifticola*, one that dwells in a rift). This MAG (S13-15_bin.75^TS^) was assembled from sequencing reads generated from samples taken from an open area located within a publicly accessible steam vent features located within the East Rift Zone. S13-15_bin.75^TS^ has a genome size of 6.8 Mb, completeness of 98.9%, and contamination of 0.4%. Both ANI and AAI identified it as a *Brasilonema* (GCA_019359415.1, ANI: 89.9%; AAI: 78.1%). In total, this MAG encodes 99 (out of 115) genes related to phycobilisome, photosynthesis, and respiratory machinery. This type genome has been deposited in ENA with the accession number ERZ28674873^TS^. Phylogenomic analysis placed this genome in the order Nostocales. *Brasilonema rifticola* belongs to the family Scytonemataceae, order Nostocales, class Cyanophyceae, and phylum Cyanobacteriota. The name has been registered under SeqCode: seqco.de/r:s5njm2s9.

#### Description of Leptodesmis pahoaensis sp. nov

*Leptodesmis pahoaensis* (pa.ho.a.en’sis (N.L. adj. *pahoaensis*), from Pāhoa, a toponym in Hawai‘i; L. suffix -*ensis*, pertaining to or originating from; N.L. adj. *pahoaensis*, originating from Pāhoa). This MAG (P23-25_bin.64^TS^) has a size of 6.8 Mb, completeness of 98.82%, and contamination of 2.2%. This MAG was assembled from sequencing reads generated from samples taken from a publicly accessible pit-like steam vent feature located within the East Rift Zone. The closest identified species is a *Leptodesmis* (GCA_037442585.1, ANI: 79.8%; GCA_021379005.1, AAI: 77.2%). In total, this MAG encodes 98 (out of 115) genes related to phycobilisome, photosynthesis, and respiratory machinery. This type genome has been deposited in ENA with the accession number ERZ28674872^TS^. Phylogenomic analysis placed this genome in the order Leptolyngbyales. *Leptodesmis pahoaensis* belongs to the family Leptolyngbyaceae, order Leptolyngbyales, class Cyanophyceae, and phylum Cyanobacteriota. The name has been registered under SeqCode: seqco.de/r:s5njm2s9.

## Discussion

### Hawaiian steam vents contain diverse, novel, and largely uncultured cyanobacteria

Bacterial community composition in steam vents can vary markedly even within closely located areas, largely due to steep gradients in temperature and moisture. Soils adjacent to active vents are typically dominated by photoautotrophic taxa (notably cyanobacteria), whereas peripheral, drier soils support drought-resistant heterotrophs, such as actinomycetes (Costello et al., 2009; Hadland et al., 2024). Previous studies report substantial diversity in the microbial assemblages of steam vent systems worldwide. In particular, the steam vents of Hawai‘i (Wall et al., 2015; Prescott et al., 2022), Paricutín and Sapichu in Mexico (Brito et al., 2019), Socompa in the Andes (Costello et al., 2009), Surtsey in Iceland (Bergsten et al., 2021), and Mutnovsky and Gorely in Russia (Allaguvatova et al., 2022), all show high cyanobacterial diversity. In contrast, cyanobacteria appear to be rare in the steam vents of Los Azufres, Mexico (Marín-Paredes et al., 2023) and Sierra Negra in the Galapagos Islands (Mayhew et al., 2007), with <1% of sequencing reads assigned to cyanobacterial sequences in Los Azufres and only a single species isolated from the Sierra Negra.

Amplicon sequencing performed in this study largely agrees with earlier studies (Wall et al., 2015; Prescott et al., 2022) that highlight the abundance of cyanobacteria in the steam vents of Hawai‘i (**Figure 1a**). Both amplicon sequencing and MAG assembly support that *Gloeobacter kilaueensis* is the dominant cyanobacterium in the pit-like environment with low-light conditions (**Figure 1b**; **Figure 2**). Leptolyngbyaceae dominate the soil and wall environments, while Hapalosiphonaceae, Oculatellaceae, Scytonemataceae, and Chroococcidiopsidaceae constitute the majority of the cyanobacterial families in the open area subjected to high-light conditions (**Figure 1c**). The diversity documented here mirrors previous findings from Hawai’i’s steam vents, including members of Gloeobacterales, Leptolyngbyales, and Nostocales (Wall et al., 2015; Prescott et al., 2022). Notably, Leptolyngbyales and Nostocales were also observed in the steam vents of Antarctica (Soo et al., 2009; Noell et al., 2025) and in Mutnovsky and Gorely, Russia (Allaguvatova et al., 2022), underscoring their wide distribution and resilience in high-temperature geothermal environments. Collectively, our findings highlight the remarkable diversity of previously undescribed cyanobacteria inhabiting Hawai‘i’s steam vents and emphasize the importance of formally characterizing this hidden diversity. The SeqCode nomenclature system greatly accelerates this process by enabling valid species descriptions in the absence of cultivation.

Here, we describe eight novel cyanobacterial species and one novel cyanobacterial genus representing five orders—Chroococcidiopsidales, Leptolyngbyales, Nostocales, Oculatellales, and Oscillatoriales—based on high-quality genome sequences and following the SeqCode guidelines (**Table 2**). Except for *Kovacikia ellensis* (W1-4_bin.70^TS^) named here, none of the new species described here had previously been sequenced, despite the availability of ∼8,000 cyanobacterial genomes in NCBI. This pattern is consistent with the hypothesis proposed by Wall et al. (2015) that steam vents act as a reservoir of yet-to-be-discovered cyanobacterial diversity.

### Recommended guideline for the description of new cyanobacterial species following SeqCode

The criteria currently used to determine whether a description of a new species applies, based on genomic data for prokaryotes—including cyanobacteria, is the 95% ANI threshold, which replaces the former 70% DNA–DNA hybridization standard (Jain et al., 2018). The same criterion is applied when determining whether a MAG represents a new species (Hedlund et al., 2022). Describing new species based on MAGs is extremely useful; however, the availability of biological material must always take priority over purely genome-based descriptions, as it allows for more applications. Therefore, in this first description of cyanobacteria under SeqCode, we propose specific guidelines that introduce three categories of biological material to refine the 95% ANI rule (**Figure 3**).

**Figure 3:**
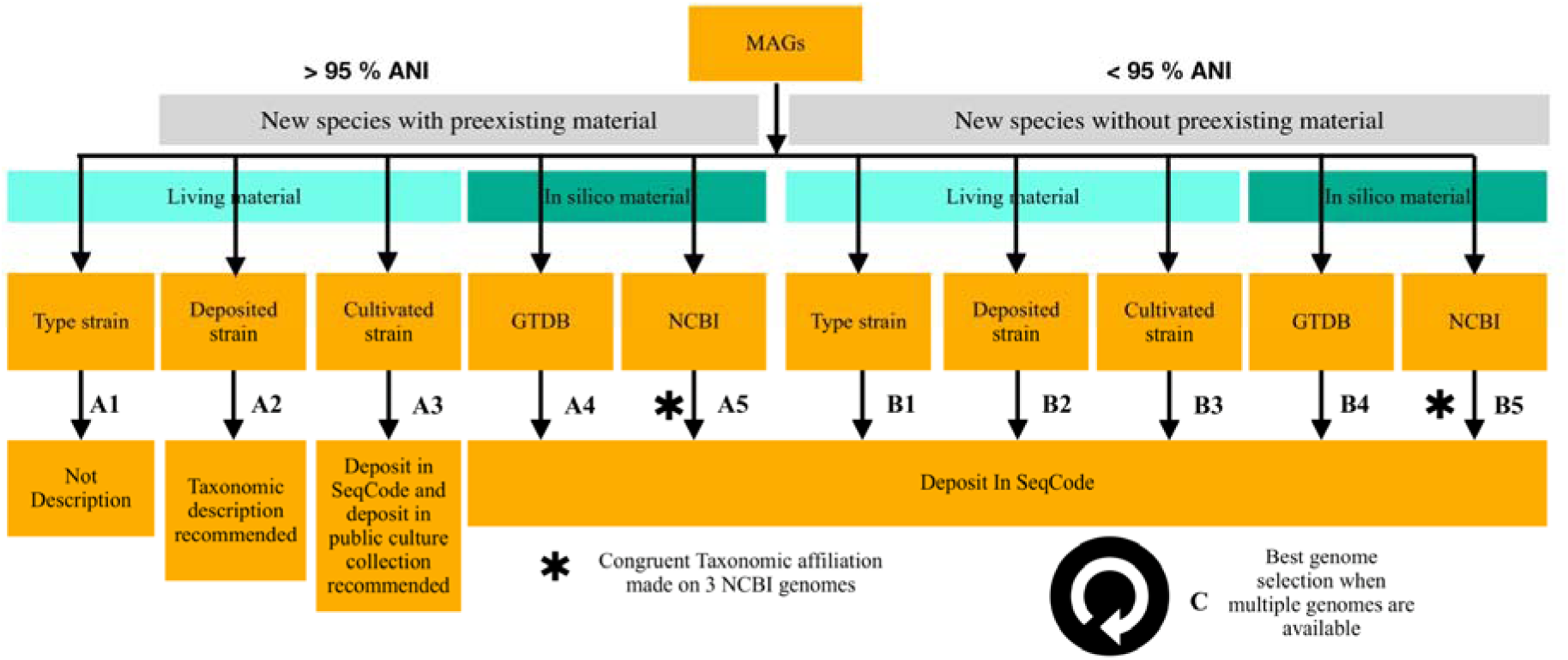
Guidelines for describing cyanobacterial species under the SeqCode. Two situations arise when determining whether a MAG can be described as a new species solely based on its genomic material, provided it meets all SeqCode quality requirements. When the MAG shares more than 95% ANI with an existing genome (Part A), different outcomes apply. If it corresponds to biological material already described as a type strain, no SeqCode registration is made since the species already exists (**A1**). If the MAG has more than 95% ANI with a strain deposited in a public culture collection, a formal taxonomic description is recommended but no SeqCode deposition occurs, giving priority to biological material (**A2**). If the MAG is over 95% identical to a cultivated strain not yet deposited in a public culture collection, a SeqCode description is performed with a recommendation to deposit the strain (**A3**), as it is unknown whether the biological material is still available. When the MAG shows more than 95% ANI with *in silico* material not yet described in SeqCode, deposition in SeqCode is recommended, either when a GTDB hit is available (**A4**) or, if not, based on at least three congruent NCBI hits (**A5**). The same logic applies when the MAG shares less than 95% ANI (**Part B**), except that hits on biological material do not imply priority and deposit in SeqCode occurs. When several genomes from the same study represent a new species, the best representative should be chosen according to SeqCode rules (**part C**).

If a MAG shares more than 95% ANI with biological material, this does not necessarily exclude a SeqCode description, as it depends on the nature of that material (**Figure 3, A part**). In the case of a MAG matching a type strain, the rule is straightforward (**Figure 3, A1**). However, two other categories can be distinguished. A MAG may share more than 95% ANI with the closest genome corresponding to a strain that has not yet been taxonomically described but is available in a public culture collection. In this case, a taxonomic description is recommended, but priority is given to the biological material (**Figure 3, A2**). A third case may occur when a MAG shares more than 95% ANI with cultivated biological material that is not deposited in a public culture collection. Since the availability of this material is uncertain, a SeqCode description can be proposed, but it should include a recommendation to deposit the strain in a public culture collection (**Figure 3, A3**), thereby linking the new genome-based taxonomy to physical biological material. This is the case for MAG W1-4_bin.70^TS^, described here as *Kovacikia ellensis*, which corresponds to a strain that has already been isolated but has not been deposited in a public culture collection.

When a MAG does not correspond to any biological material but instead matches *in silico* data at over 95% ANI, and those data are not yet described in SeqCode, a genome-based species description applies (**Figure 3, A4–A5**). A special case arises when a MAG lacks any GTDB match but has hits only in the NCBI database. Because the GTDB taxonomy is unavailable in this scenario, classification relies solely on the less curated NCBI taxonomy (Parks et al., 2022). We recommend ensuring congruence by considering the last common ancestor (LCA) of the three closest NCBI hits. An example is provided for three of our samples (HW-12_bin.63, S2-4_bin.10, and S29-31_bin.283), where the taxonomy of the three top NCBI hits differed too greatly to allow a SeqCode description (**Table S3**). When a MAG shares less than 95% ANI, it falls under the conventional framework commonly used for describing prokaryotic species (**Figure 3**).

In addition to species descriptions based on ANI, an AAI threshold of 65% should be used to determine the presence of a novel genus, which was not the case in this study. However, as illustrated by MAG W1-4_bin.36^TS^, taxonomy of closely related ANI matches does not always allow for a clear genus assignment, since the closest genomes are not classified with sufficient precision, here at the Nostocales order level. In such cases, phylogenetic placement provides the necessary resolution, as exemplified here by the establishment of the new genus *Paranostoc speluncae*.

## Contributions

ES performed the metagenomic analyses, BD and MH carried out the amplicon sequencing analyses, LC established the guidelines for taxonomic description. ES, BD, LC, and JS wrote the manuscript. All authors read and approved the final version of the manuscript.

## Acknowledgments

This work was supported by a research grant (PDR T.0018.24 OR-OX-PHOT-IN-CYN) from the Belgian National Fund for Scientific Research (F.R.S.-FNRS) to DB. LC is supported by a mandate from the Belgian National Fund for Scientific Research (F.R.S.-FNRS). JS is supported by startup funds from the George Washington University and a US National Science Foundation (NSF) grant (award number 2442122).

## Supplementary Table Legends

**Supplementary Table 1.** Relative abundance (%) of the taxa, at different taxonomic levels (Phylum, Family and Genus) in the four sampling sites.

**Supplementary Table 2.** Statistics and taxonomic classifications of 40 high-quality cyanobacterial and melainabacterial MAG assemblies

**Supplementary Table 3.** The number of detected gene sequences in the nine newly characterized MAGs, categorized by functions related to the phycobilisome, photosynthesis, and respiratory machinery.

